# Natural selection shapes variation in genome-wide recombination rate in *Drosophila pseudoobscura*

**DOI:** 10.1101/787382

**Authors:** Kieran Samuk, Brenda Manzano-Winkler, Kathryn R. Ritz, Mohamed A.F. Noor

## Abstract

While recombination is widely recognized to be a key modulator of numerous evolutionary phenomena, we have a poor understanding of how recombination rate itself varies and evolves within a species. Here, we performed a comprehensive study of recombination rate (rate of meiotic crossing over) in two natural populations of *Drosophila pseudoobscura* from Utah and Arizona, USA. We used an amplicon sequencing approach to obtain high-quality genotypes in approximately 8000 individual backcrossed offspring (17 mapping populations with roughly 530 individuals each), for which we then quantified crossovers. Interestingly, variation in recombination rate within and between populations largely manifested as differences in genome-wide recombination rate rather than remodeling of the local recombination landscape. Comparing populations, we discovered individuals from the Utah population displayed on average 8% higher crossover rates than the Arizona population, a statistically significant difference. Using a Q_ST_-F_ST_ analysis, we found that this difference in crossover rate was dramatically higher than expected under neutrality, indicating that this difference may have been driven by natural selection. Finally, using a combination of short and long read whole-genome sequencing, we found no significant association between crossover rate and structural variation at the 200-400kb scale. Our results demonstrate that (1) there is abundant variation in genome-wide crossover rate in natural populations (2) interpopulation differences in recombination rate may be the result of local adaptation, and (3) the observed variation among individuals in recombination rate is primarily driven by global regulators of crossover rate, with little detected variation in recombination rate among strains across specific tracts of individual chromosomes.

## Introduction

Meiotic recombination is the exchange of genetic material between homologous chromosome that occurs during meiosis. This exchange takes two forms, crossing over and non-crossover gene conversion, both of which are initiated by the formation of a double-strand break during meiosis. Recombination, particularly crossing over, appears to be an important mediator of chromosome pairing during meiosis, with most species exhibiting an average of one crossover per chromosome arm (Dapper & Payseur, 2017; Hughes *et al*., 2018).

While physical constraints often set a lower bound on rates of recombination, the evolution of recombination rate and particularly the rate of crossing over (i.e., number of crossovers per generation in a genomic interval) can have far-reaching effects on nearly every evolutionary process (Charlesworth *et al*., 2009; Dapper & Payseur, 2017; Ritz *et al*., 2017). For example, recombination rates can modulate processes as diverse as adaptation to a new environment, the evolution of reproductive isolation, and the dynamics of introgression between populations (Barton & Otto, 1997; Otto & Lenormand, 2000; Marais & Charlesworth, 2003; Martin *et al*., 2019). More generally, recombination rate determines the degree to which an individual’s parental chromosomes are mixed in their gametes – i.e., how often novel allelic combinations are generated in their gametes. Increases or decreases in this rate can be favored under different evolutionary/ecological conditions. For example, increasing the rate of recombination can facilitate adaptation by increasing the probability that adaptive and maladaptive alleles will be decoupled or that adaptive alleles will be brought together in the same genotype (i.e., overcome Hill-Robertson interference (Barton, 2010). Increased rates of recombination are similarly favored when fitness optima change rapidly between generations, e.g., under fluctuating selection (Charlesworth, 1976). On the other hand, lower recombination rates can be favored under scenarios in which adaptive combinations of alleles are at risk of being broken apart, such as under maladaptive gene flow (Kirkpatrick & Barton, 2006). Reduction/suppression also appears to have important consequences for the evolution of reproductive isolation (Noor *et al*., 2001; Kirkpatrick & Barton, 2006) and patterns of introgression and divergence in the genome (Samuk *et al*., 2017; Schumer *et al*., 2018; Martin *et al*., 2019).

While there is a rich theoretical literature focused on the evolution of recombination rate, empirical studies have lagged somewhat behind. One reason for this may be that recombination rate is difficult to quantify directly – it generally requires the construction of a linkage map from a genetic cross and/or cytological visualization of recombination-associated proteins (Kulathinal *et al*., 2008; Dapper & Payseur, 2017; Wang & Payseur, 2017). Recently, many studies have attempted to overcome this difficulty by instead estimating a population genetic quantity known as ρ, the population scaled recombination rate (Stumpf & McVean, 2003). This quantity is the product of four times the effective population size (*N*_*e*_) and realized recombination rate (sometimes denoted “c”) (Wall, 2000). The general approach to estimating ρ is to perform coalescent simulations and fit a simulated value of ρ to observed patterns of linkage disequilibrium (LD) (Auton & McVean, 2007; Gärtner & Futschik, 2016; Adrion *et al*., 2019). While this approach has proven successful at recapitulating many of the general features of the recombination landscape in many species, it is not able to disentangle changes in *N*_*e*_ or LD *per se* (e.g. as a result of selection or demography) from changes in recombination rate (either locally or genome-wide) (Nei & Li, 1973; Adrion *et al*., 2019). Further, these methods are highly sensitive to increases in LD that occur as a result of gene flow between populations (Nei & Li, 1973; McVean, 2007; Cutter, 2019). As such, LD-based methods are likely to be less appropriate for the study of the evolution of recombination rate than direct estimates of recombination rate.

In spite of methodological difficulties, there has been a recent resurgence of interest in the empirical study of the evolutionary causes and consequences of recombination rate (Dapper & Payseur, 2017; Ritz *et al*., 2017; Stapley *et al*., 2017). One key contributor to this resurgence has been the democratization of high throughput genotyping, which has increased the tractability of creating high density linkage maps in non-model species (e.g. using pedigreed populations or gametic sequencing, (Hellsten *et al*., 2013; Johnston *et al*., 2016). The increased availability of such linkage maps has in turn led to a growing appreciation of the enormous diversity in recombination rate that exists between taxa (Stapley *et al*., 2017). This variation can manifest globally, i.e. genome-wide, or locally, i.e. along a specific tract of a chromosome (Smukowski & Noor, 2011; Stapley *et al*., 2017).

Studies using direct estimates of recombination rate have largely focused on describing differences in recombination between species or sexes (Dumont & Payseur, 2008; Hunter *et al*., 2016; Stapley *et al*., 2017). However, there are surprisingly few studies focused on directly testing evolutionary hypotheses concerning variation in recombination rate. For example, a key question that emerges from the theoretical literature is: what role does natural selection play in shaping rates of recombination (Charlesworth, 1976; Barton & Otto, 1997; Otto & Michalakis, 1998)? In other words, does natural selection shape recombination rates, perhaps as result of environment or mutational load? While a tempting research direction, testing such an adaptive hypothesis is non-trivial with traits that are difficult to measure and manipulate such as recombination rate (Dapper & Payseur, 2017; Ritz *et al*., 2017). One approach may be experimental evolution, in which the proposed selective agent that favors/disfavors changes in recombination rate is experimentally varied, evolved differences in recombination rate are quantified, and these differences are then compared to a null (non-adaptive) expectation (Aggarwal *et al*., 2015). This approach is powerful but highly laborious and difficult to apply to natural systems. A more broadly applicable method for detecting the influence of natural selection a quantitative trait is perhaps the Q_ST_-F_ST_ approach (Leinonen *et al*., 2013). Originating in the quantitative genetics literature, this powerful method is designed to answer the question: are the observed differences between populations in a quantitative trait greater than expected on the basis of drift alone (Whitlock, 2008; Leinonen *et al*., 2013)? This question is formalized as a statistical hypothesis test that compares variation in a quantitative trait (Q_ST_) within and between populations to a null distribution of variation in neutral genetic markers (F_ST_) within and between populations (Whitlock, 2008; Whitlock & Guillaume, 2009). While the Q_ST_-F_ST_ method has enjoyed great success in the quantitative and evolutionary genetics literature, it has not yet been applied to testing the role of selection on recombination rate. Given its flexibility and applicability to any quantitative trait, we perceive Q_ST_-F_ST_ an ideal method for exploring the evolution of recombination rate.

Along with quantifying intraspecific variation and the role of natural selection, we also have a poor understanding of the genetic basis differences in recombination rate between populations and species. As is the case for other traits, identifying the genetic architecture of evolutionary changes in recombination rate allows for a more complete explanation for how and why recombination rate evolves (Barrett & Hoekstra, 2011). One specific question is the degree to which variation in recombination rate manifests as a local vs. global phenomenon. Local variation in recombination can arise due to structural variants that suppress recombination such as inversions and large deletions (Gaut *et al*., 2007; Völker *et al*., 2010; Morgan *et al*., 2017). In contrast, global variation can arise from mutations in the genes involved in meiosis and/or double-strand break repair pathways (Brand *et al*., 2018). Modifiers of both global and local rates of recombination have been identified in laboratory and/or interspecific crosses, but their occurrence in natural populations of individual species is only just beginning to be explored (Hunter *et al*., 2016; Brand *et al*., 2018). Notably, most theoretical studies of recombination rate have focused on variation in global rates of recombination, e.g. via ‘modifier’ models, but it is unknown if such modifiers are common in natural populations.

Here, we performed a comprehensive study of recombination rate (meiotic rates of crossover) in two natural populations of *Drosophila pseudoobscura* from Utah and Arizona, USA. We made use of modern sequencing and genetic map construction methods, along with the Q_ST_-F_ST_ approach. The key questions we sought to answer were:

(1) How does recombination rate vary within and between natural populations of *D*. *pseudoobscura?*

(2) Are the interpopulation differences in recombination the result of neutral drift or natural selection?

(3) Do genetically-based differences in recombination rate manifest largely as local variation or global variation? To what degree are these differences connected to structural variation and allelic variation in recombination-associated genes?

By answering these three questions, we hope to extend our understanding of the evolution of recombination rate in *D*. *pseudoobscura* and provide proof-of-concept of the tractability of studying the evolutionary genetics of recombination rate in natural populations.

## Methods

### Collection and inbreeding of wild samples

We collected wild *D*. *pseudoobscura* from Madera Canyon, AZ, USA (31°42’48.9”N, 110°52’22.4”W) and American Fork Canyon, UT, USA (40°26’38.9”N, 111°42’08.5”W) in May and July of 2015 respectively using bucket traps (Ritz & Noor, 2015). These populations were chosen because they were known to share similar karyotypic configurations (e.g. inversions) but also differ in their ecological context (i.e. xeric vs. sub-alpine). We returned live individuals to the laboratory, isolated females, and created inbred lines from their offspring (one line per surviving female). These lines were created by successive crosses between virgin siblings for a minimum of 14 generations. The inbred lines (and all subsequent lines) were reared in 20C incubators with 65% relative humidity and photoperiods of 14D:10N. The inbreeding process resulted in a total of 7 inbred lines from Arizona and 12 from Utah.

### RAD-seq libraries from wild samples

To generate a set of SNPs for estimating F_ST_ between the Utah and Arizona populations, we performed double-digest RAD-seq reduced representation sequencing. To begin, we extracted DNA from single wild-caught individuals (excluding the females used to initiate the inbred lines) via phenol-chloroform DNA extraction. We then performed a RAD-seq library preparation protocol after (Peterson *et al*., 2012). The resulting libraries were sequenced in a single lane on an Illumina HiSeq 4000 at the Duke Center for Genomic and Computational Biology sequencing facility.

### Whole genome sequencing of inbred lines

We performed both short read and long read whole genome sequencing on all 19 inbred lines, as well as our testers line (MV2-25 and Flagstaff-14). The short read libraries were prepared by first performing phenol-chloroform DNA extractions from pools of 20-30 individual female flies. We quantified DNA purity and concentration via Nanodrop (Thermofisher Inc.) and Qubit (Qiagen Inc.). The DNA samples were then submitted for library preparation and sequencing via Illumina NovaSeq (300-400bp insert, 150bp paired end reads) at the Duke Center for Genomic and Computational Biology sequencing facility.

The long read libraries were prepared by first performing high-molecular weight DNA extractions from pools of 20-30 female flies using Qiagen Midi/Mini Prep DNA extraction kits (Qiagen Inc.). These were then assessed for fragment size via standard gel electrophoresis, and submitted for sequencing on a PacBio Sequel (4 SMRT cells, 4-5 samples multiplexed per cell) at the Duke Center for Genomic and Computational Biology sequencing facility.

### Whole genome variant calling: short read WGS and RAD-seq data

We identified variants in the short read data (both isoline whole genome sequencing and wild population RAD-seq) using an analysis pipeline based on the GATK best practices (DePristo *et al*., 2011; Van der Auwera *et al*., 2013). The complete code for this pipeline is available as a Github repository at http://github.com/ksamuk/recomb_evo (available pre-publication upon request). All tools were run with default settings unless otherwise indicated. Briefly, we aligned the reads for each sample to the *D*. *pseudoobscura* reference genome (version 3.04 from FlyBase, ftp://ftp.flybase.net/genomes/Drosophila_pseudoobscura/) using bwa mem version 0.7.17 (Li & Durbin, 2009). We marked adapters and duplicates using PicardTools (Wysoker *et al*., 2013), and performed individual-level genotyping for each set of marked reads using the HaplotypeCaller. We then performed joint genotyping on the resulting set of GVCFs via GenotypeGVCFs. We filtered SNPs in the resulting VCF using the GATK Best Practices hard filters (see scripts for details), working in R 3.4.1 (R Core Team, 2018) with the vcfR and tidyverse packages (Knaus & Grünwald, 2017; Wickham, 2017).

### Creation of mapping populations

To estimate variation in crossover rate in our inbred lines, we created backcross-like mapping populations (crossing scheme shown in Figure 1). We crossed groups of 3-5 males from each isoline to single virgin females from the *D*. *pseudoobscura* reference genome isoline (MV2-25, provided by Dr. Steve Schaeffer). We then allowed the F_1_ offspring to develop, and collected virgin females from the resulting offspring. Finally, we crossed these virgin F_1_ females to males from a second fixed isoline, Flagstaff-14 (a highly inbred isoline from Flagstaff, AZ). This resulted in a backcross-like mapping population for each of the AZ and UT lines, in which all BC1 offspring had one maternal chromosome from their F_1_ mother and one paternal Flagstaff-14 chromosome with a fixed, known genotype (Figure 1A). This design allows for straightforward mapping of recombination events that occurred in F_1_ females. As such, our estimates are unable to detect any variation in recombination due to recessive-acting effects and may underestimate total recombination rates (e.g. from modifiers that act in an additive fashion) in the pure inbred lines. Critically, this potential underestimation is identical across all F_1_ families, and thus cannot (in and of itself) generate systematic differences in recombination rate between lines or populations.

**Figure 1.**
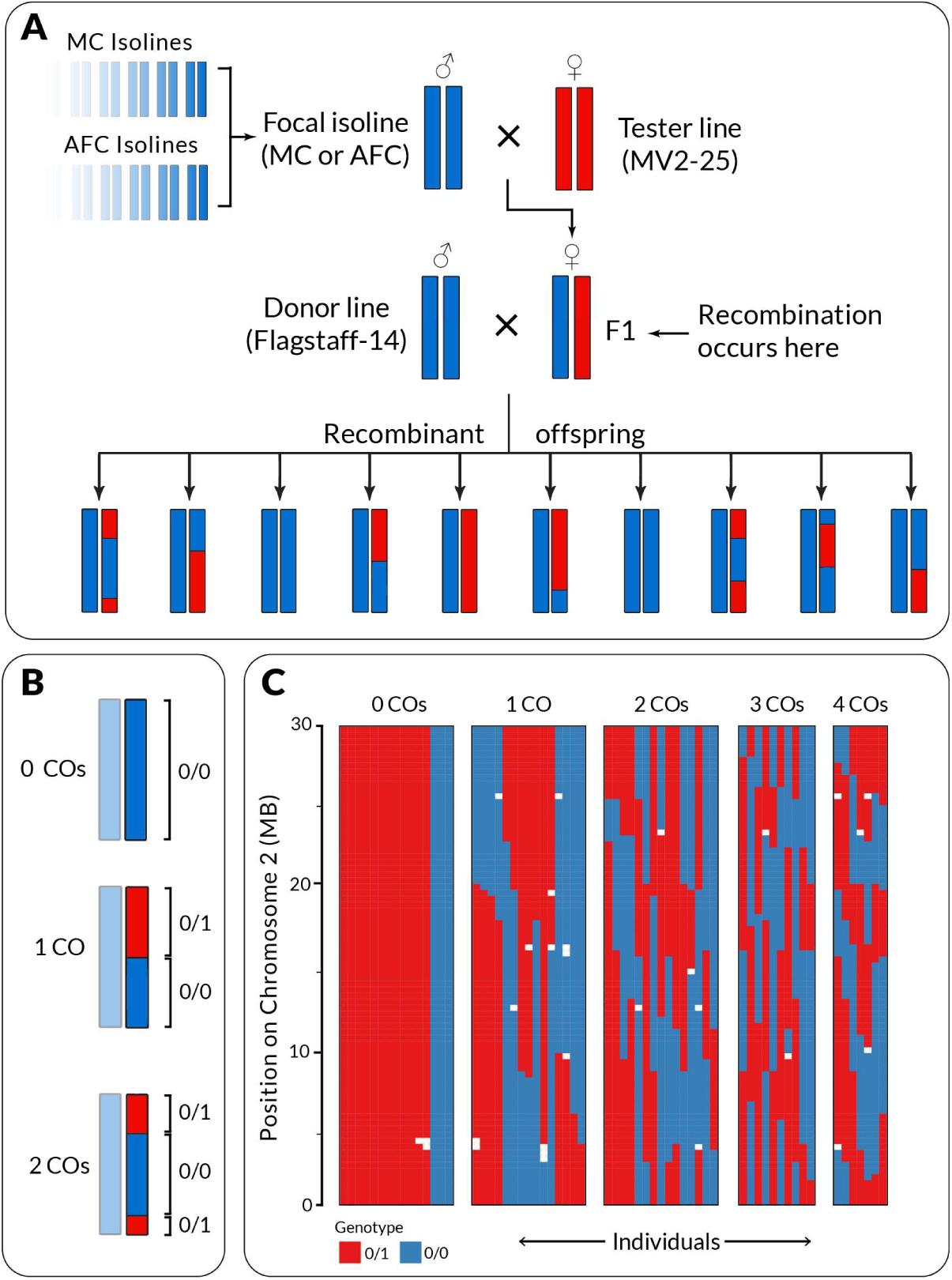
Schematic of the crossing design and one method of interfering crossovers. (a) Isolines from MC and AFC were individually crossed to tester lines to generate F1s, which were subsequently crossed to a “donor line” sharing the same genotype as all isolines, but a different genotype than the tester line at all marker loci. Further, all markers were selected such that only two alleles were found in all lines, with the tester line having one allele (“1”) and all other lines including the donor line having the other (“0”). This allows for the scoring of crossovers as changes in heterozygosity, as shown in (b). (c) Example genotypic data from one chromosome showing the number of inferred crossovers. White genotype states indicate missing data.

### Genotyping of mapping populations

Because our goal was to quantify the number of crossovers per generation rather than their precise location, we performed low density, genome-wide SNP genotyping using an amplicon sequencing approach. To do this, we adapted the ‘GT-seq’ method outlined in (Campbell *et al*., 2015). To begin, we identified SNPs genotyped in the whole genome dataset that were unique to the MV2-25 isoline (i.e. fixed for one allele in all 19 inbred lines and Flagtaff-14 and fixed for another allele in MV2-25). Genotyping these markers in BC1 individuals allows the recovery of genotypic phase simply by examining the genotype of the marker SNPs -- regions with UT or AZ ancestry are represented as runs of heterozygous SNPs and regions with MV2-25 ancestry are represented as runs of homozygous SNPs (see diagram in Figure 1B). In total, we selected 500 of these SNPs evenly spaced at approximately 300kb intervals along each chromosome.

We designed primer pairs to generate ∼200-300BP amplicons containing each of our target SNPs. These primer pairs were optimized to minimize primer-primer interactions during multiplex PCR (primer design service provided by GT-Seek Ltd., Idaho, USA). With these primers in hand, we performed two test library preps using the GT-seq protocol described in Campbell et al. (2015). We sequenced the first test library on a MiSeq (V3 flowcell, Illumina Corp., California, USA), and identified poorly performing amplicons using the criteria outlined in Campbell et al. (2015), i.e. high dropout, low representation among individuals, evidence of amplicons mapping to duplicate regions, etc. (service provided by GT-Seek LTD, Idaho, USA). We then prepared a second test library with the primers for the poor-performing amplicons omitted and sequenced it as above. A final screen for poor-performing amplicons resulted in a final set of 390 amplicons ranging from 200-300bp, each containing at least one recombination-informative SNP.

After optimizing our panel of amplicons, we used GT-seq to genotype approximately 400 BC1 offspring from each mapping population (400 individuals from each of 19 lines, a total of approximately 7600 individuals). We created two pools of 40 plates (individuals and plates are individually barcoded as part of GT-seq library preparation) and submitted these for sequencing on an Illumina NovaSeq (Mid output flow cell, Illumina Corp., California) at the Duke Center for Genomic and Computational Biology sequencing facility. We called SNPs in our sequenced GT-seq amplicons using an identical approach to our whole genome short read data. The final dataset contained 420 total variants across all amplicons, sequenced to an average depth of ∼200X. We performed further quality control on the resulting SNPs in R using the vcfR and tidyverse packages (Knaus & Grünwald, 2017; Wickham, 2017). First, we dropped any markers that mapped to genomic locations outside our original targeted amplicons. Next, we dropped any individuals that had an average depth below 10x (19/7600 individuals). Finally, we removed any markers that displayed any evidence that they were in fact not unique to the tester line. This was done by removing markers displaying: (1) any evidence of segregation distortion, (2) any evidence that any of the isolines were in fact polymorphic for the marker or (3) high dropout (i.e. represented in fewer than 75% of samples). In some cases, the source of marker dropout was clearly an undetected INDEL polymorphism in the amplified regions, which, for consistency among lines, we erred on the side of removing rather than recoding as them as markers for mapping. After filtering, we recoded all SNP genotypes as 0 for the isoline/donor line state and 1 for the tester line state. Because of the backcross design, the only possible genotypes were thus 0/0 and 0/1.

### Detection of recombination events

We identified crossovers in two steps: (1) ancestry assignment of chromosome segments and (2) crossover counting. To begin, we updated the genomic ordering of our markers using the genomic scaffold ordering from Schaeffer *et al*. (2008). After markers had been reordered, we assigned the ancestry (isoline or tester) of chromosomal segments by identifying runs of 0/0s and 0/1s. In regions with a single ancestry assignment, we imputed across gaps of missing markers (e.g. due to filtration or dropout) shorter than 2 markers (∼400kb). After local ancestry was assigned, we counted crossovers by counting the number of ancestry changes (from 0/0 to 0/1) along each chromosome in each individual. Following (Broman & Sen, 2009), we ignored double crossovers spanning less than 2 markers (400kb) (crossover interference should make close range double crossovers exceedingly rare, and thus these cases likely represent genotyping or marker-order errors).

This crossover counting method assumes that the order of markers on each chromosome is identical in each line. Differences in marker order could, for example, generate spurious double crossovers (although ignoring short double crossovers somewhat ameliorates this issues). To directly address the possibility of different marker orders among lines, we created separate genetic maps for each isoline using the R packages r/QTL and ASMap (Broman & Sen, 2009; Taylor & Butler, 2017). Following the general recommendations from the documentation, these two packages agnostically infer linkage group assignment, marker order, and genetic distances between markers. Overall, there was high concordance in marker order between all the individually-inferred maps (Figure S1, Figure S2). Individual recombination rate estimates within each line were highly similar when using the reference genome marker order or individually-inferred marker orders (Figure S2, Spearman rank correlation = 0.93, p < 2.2×10^−16^). We thus elected to use the reference genome marker order for all subsequent analyses.

### *Q*_*ST*_-*F*_*ST*_ analysis

To test the hypothesis that population-level differences in recombination rates are driven by natural selection, we performed a *Q*_*ST*_ -*F*_*ST*_ analysis (Whitlock & Guillaume, 2009; Leinonen *et al*., 2013; Walsh & Lynch, 2018). We began by computing a point estimate of *Q*_*ST*_ for genome-wide recombination rate using the R package lme4 by fitting a linear mixed effects model with the following form: crossover count = intercept + (1|line) + (1|population). We extracted the variance components for population and family:population using the R function varcomp(). Following Walsh & Lynch (2018) we computed *Q* _*ST*_ using the following formula:

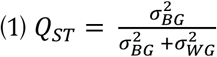

Where 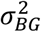 denotes the between-group variance and 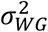 denotes within-group variance. Note that the within-group (i.e. between inbred line) variance term in the denominator is not multiplied by two in the case of haploids or completely inbred lines (Walsh & Lynch, 2018).

We computed *F*_*ST*_ using SNPs genotyped via RAD-seq in wild AZ and UT individuals. To do this, we converted the GATK VCF to a SNP table using vcfR and the tidyverse package in R (see analysis scripts). We then converted the resulting SNP table for manipulation in the R package SNPRelate (Shen *et al*., 2012). Using SNPRelate, we first performed LD pruning (default settings, r < 0.2) to reduce statistical non-independence between SNPs (Lotterhos, 2019). This resulted in a dataset composed of 16 individuals for AZ and 42 for UT, with a total of 6 591 high quality SNPs. We then computed per-SNP estimates of Weir and Cockerham’s *F*_*ST*_ using SNPRelate, requiring filtered sites to have a minimum minor allele frequency of 0.1.

We assessed the statistical departure from neutrality for each value of Q_ST_ using the Null-QST method outlined in Whitlock & Guillame (2009) and Walsh & Lynch (2018), with a modification to accommodate trait data from inbred lines. The general approach outlined in these two references is to simulate the expected distribution of *Q*_*ST*_ for a neutral trait (denoted 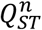, neutral Q_ST_) via a parametric bootstrap, and use this distribution as the basis of a statistical test of the hypothesis 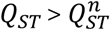.

To simulate the distribution of 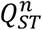 we first estimated the between-group 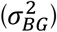 and within-group 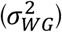 variance components. We obtained these values via REML estimation by fitting mixed-effects linear models using the function lmer in the R package lme4 (Bates *et al*., 2007). These models took the form *crossovers ∼ intercept + (1/population) + (1/line)*. We extracted the variance components (standard deviations of the random effects) using the function VarComp from lme4. We next generated 10 000 (nonparametric) bootstrap estimates of the mean value of Weir and Cockerham’s *F*_*ST*_ by resampling the RADseq SNPs with replacement, and computing genome-wide mean *F*_*ST*_ using SNPRelate. We then generated 10 000 matching parametric bootstrap estimates of the 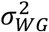 by multiplying the REML point estimate by a random draw from a *χ*^2^ distribution with degrees of freedom equal to the number of inbred families (df = 17). Next, we generated parametric bootstrap estimates of the expected values of 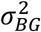 for a neutrally evolving trait using the equation:

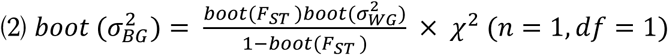

Modified from Whitlock and Guillaume (2009) to accommodate complete inbreeding. In equation (2), “boot” indicates individual bootstrap samples for each quantity, and the *χ*^2^ term represents a draw from a *χ* ^2^distribution with degrees of freedom equal to the number of populations minus one (one, in this case). This procedure results in 10 000 bootstrap samples for 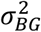 and 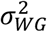, from which we computed 10 000 bootstrap samples of 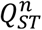 using equation (1). We finally computed a p-value for the observed value of *Q*_*ST*_ by determining the number of 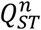 values that exceeded the observed value of *Q*_*ST*_ distribution. We also computed a confidence interval for *Q*_*ST*_ − *F*_*ST*_ (the difference between *Q*_*ST*_ and *F*_*ST*_, expected to be zero under the neutral model) by subtracting each value the 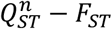 distribution from the observed value of *Q*_*ST*_ − *F*_*ST*_ (after Walsh & Lynch 2018).

### Candidate genes associated with recombination differences

We explored the possibility that between-line variation in meiosis-related candidate genes may underlie between-line differences in recombination rate. We were specifically interested in the possibility that coding changes in meiosis genes might underlie differences in recombination rate. To do this, we first assembled a list of candidate genes from Anderson *et al*. 2009 and Hunter *et al*. 2016 (Anderson *et al*., 2009; Hunter *et al*., 2016). We then obtained the FASTA sequences for these genes in each line by intersecting the short read variant calls (including INDELs) with the *D*. *pseudoobscura* reference genome CDS for each candidate gene. To ensure proper alignment, we then performed multiple alignment of the line-level FASTA sequences and the reference CDS using MAFFT version 7.407 (Katoh & Standley, 2013). Once the sequences had been aligned, we identified non-synonymous, non-reference alleles in each line. We then tested for associations between line-level recombination estimates and genotype at each site where at least one non-synonymous change occurred in each gene. To this end, we fit linear models with recombination rate as the response and genotype (at all variable non-synonymous sites) as the predictor. This yielded a p-value for each genotype vs. recombination comparison, which we adjusted via the FDR approach (Benjamini & Hochberg, 1995), with FDR < 0.05 adjustments performed using the function p.adjust in R.

### Association between local structural variation and recombination rate

Along with the candidate gene approach to examine associations with genome-wide recombination rate, we also investigated the possibility that small-scale differences in genomic structure between the inbred lines may explain differences in recombination rate. This may be of particular importance given that our design required measuring recombination rate in F_1_ individuals (inbred line × tester line), and that structural heterozygosity has a well-known negative association with recombination rate (Morgan *et al*., 2017; Crown *et al*., 2018; Korunes & Noor, 2019).

To test if differences in genome structure underlie local differences in recombination rate in our inbred lines, we first identified structural variants (SVs) using two approaches. First, we used the SVtools pipeline (Larson *et al*., 2019)to identify SVs using paired-end short read data. This pipeline identifies structural variation using a variety of genomic signatures, particularly split reads (different parts of a single read mapping to multiple discrete locations) and discordant reads (paired end reads separated by a much greater genomic distance than expected on the basis of their insert size). The general procedure is to identify split/discordant reads using the tools *SAMBAMBA* and *SAMBLASTER*, which are then analyzed and annotated with the SVtools variant callers (Faust & Hall, 2014; Tarasov *et al*., 2015). The resulting structural variant VCF was filtered via empirical cut offs using the guidelines in (Larson *et al*., 2019). Along with SVtools, we separately identified structural variation in the PacBio long reads dataset using the PacBio structural variant pipeline and tools, *pbsv* (https://github.com/PacificBiosciences/pbsv, see also Wenger *et al*. (2019). This involves aligning the long reads with *minimap2* (accessed via the *pbmm2* wrapper), identifying individual signatures of structural variation using pbsv, and jointly calling structural variation from the combined set of signatures. This again results in a VCF containing structural variants, which we filtered using empirical cutoffs as before.

After identifying structural variants, we next quantified the total difference in sequence homology between each line and the tester line (MV2-25) for each genomic interval where recombination was measured (∼300kb windows). To do this, we summed the total number of non-shared, non-reference base pairs between each line and the tester line. We included SNPs, inversions, insertions, deletions, and translocations in this calculation. This method collapses multiple classes of genomic variation into a single, consistent metric and avoids the ambiguity associated with identifying shared locations of breakpoints for the structural variants (e.g. needed for per-variant associations). Further, this method focuses on the most likely biological cause of structurally-mediated recombination suppression, i.e. differences in homology *per se*, which has been widely demonstrated in many species (Greig *et al*., 2003; Salomé *et al*., 2012; Rogers *et al*., 2018). We also tabulated the total count of structural variant alleles (of any type) that differed between each isoline and the tester line for each recombination interval. We normalized all homology estimates and structural variant counts in each window using both the total number of genotyped base pairs in each window as well as the mean depth per isoline.

We tested the association between normalized sequence homology and recombination rate via a hierarchical linear model fit using the function glmer from the R package lme4 (Bates *et al*., 2007). This model had the form: *recombination rate* ∼ *method* * *homology* + (*1/window identity*) + (*1/inbred line*), with Poisson-distributed errors and a log link function. Assigning window identity (i.e. genomic region in which recombination and homology were measured) as a random effect controls for mean local variation in recombination rate (i.e. it normalizes the absolute recombination rates among windows). Similarly, modelling inbred line identity as a random effect controls for genome-wide differences in recombination rate, which are unrelated to local variation. We assessed the significance of the homology term in the model by comparing the full model to a model with only random effects via a likelihood ratio test in R. We finally repeated this model fitting procedure with the normalized count of differing structural variant alleles as the predictor.

## Results

### Within and between population variation in recombination rate

Genome-wide recombination rate varied significantly within and between the D. pseudoobscura populations we studied. Within lines, there was a range of 3.75–4.75 crossovers per genome, corresponding to 0.75-0.95 crossovers per chromosome arm on average (Figure 2A). This between-line variation was statistically significant (p < 2.2×10^−16^, Likelihood Ratio Test Statistic = 141.13, df = 1, comparison via dropping inbred line random effect). At the population level, lines from American Fork Canyon, UT had 5.20 ± 0.17 crossovers per genome on average, while lines from Madera Canyon, AZ had 4.82 ± 0.21 crossovers per genome on average, a significant difference in genome-wide crossover rate (Figure 2B, Type II Wald *χ*^2^ Test, p = 0.018, df = 1).

**Figure 2.**
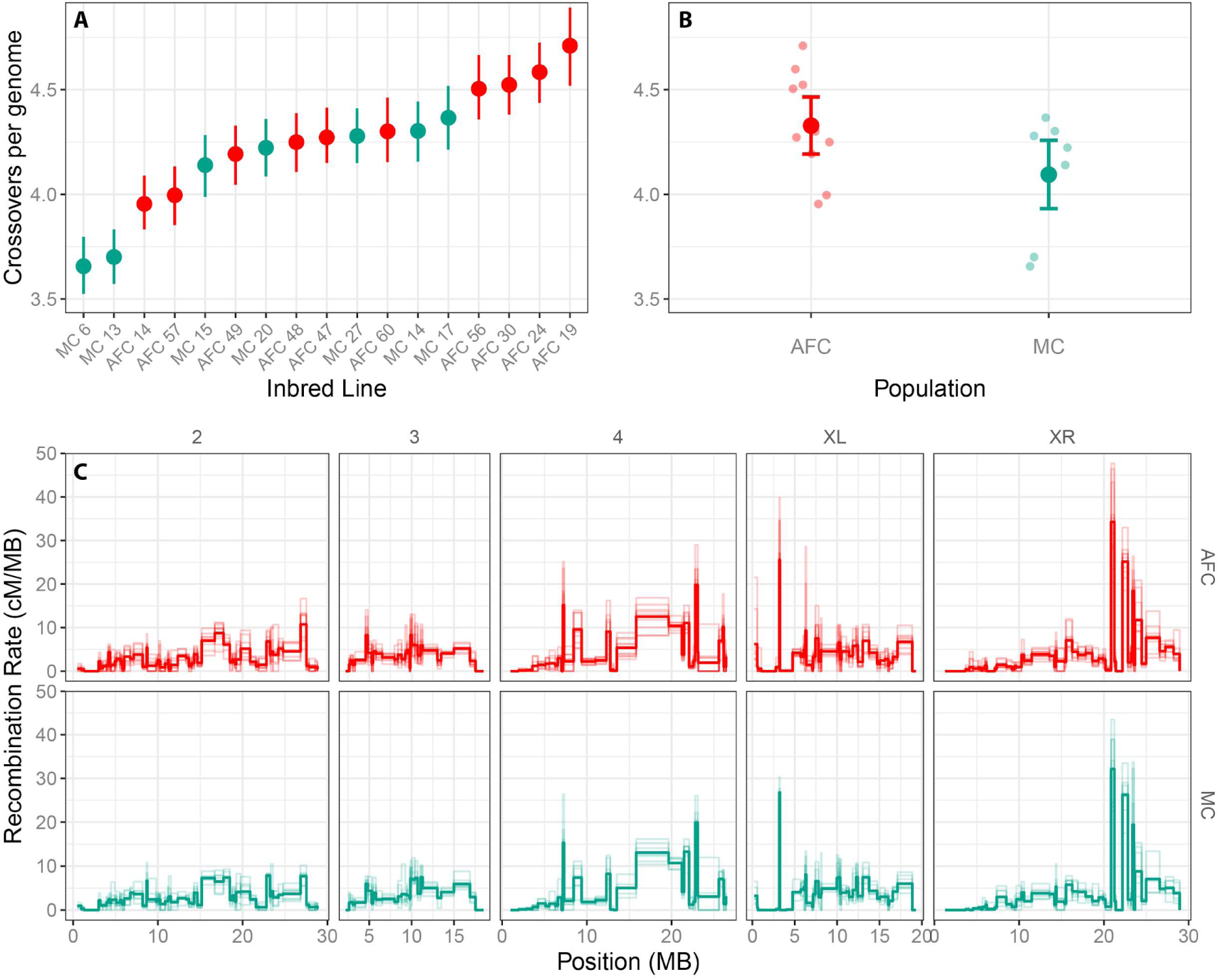
Recombination rate varies within and between populations of *D*. *pseudoobscura*. (A) Variation in genome-wide crossing over frequency for 17 inbred lines. Lines are colored according to their population of origin (Green, MC: Madera Canyon, AZ, Red, AFC: American Fork Canyon, UT.). Points depict the mean crossover frequency for each line with vertical lines representing 95% confidence intervals (n = 384 per line). (B) Differences in crossover frequency between AFC and MC. Jittered points are individual line means (from A), and larger points are marginal means derived from mixed model regression coefficients along with 95% confidence intervals (error bars). (C) Variation in recombination rate across the genome. Each panel depicts recombination rate along a single chromosome arm (columns) in one of two populations (rows). Thick lines depict population average recombination rates, with lighter lines depicting rates for individual inbred lines.

That said, despite genome-wide differences, the structure of variation in recombination rate along the genome was extremely similar among individuals and populations (Figure 2C & Figure 3A, R^2^ = 0.96, correlation test t = 68.866, df = 207, < 2.2×10^−16^). Interestingly, we also found that chromosome-scale recombination rates were highly correlated *within* lines, such that there was a strong trend that lines with high recombination rate on one chromosome tended to also have high recombination on other chromosomes (Figure 3B; average R^2^ = 0.78, all correlations significant via correlation tests, p <0.0001). In sum, these results suggest that most phenotypic variation in recombination rate within and between populations occurs at the genome-wide scale.

**Figure 3.**
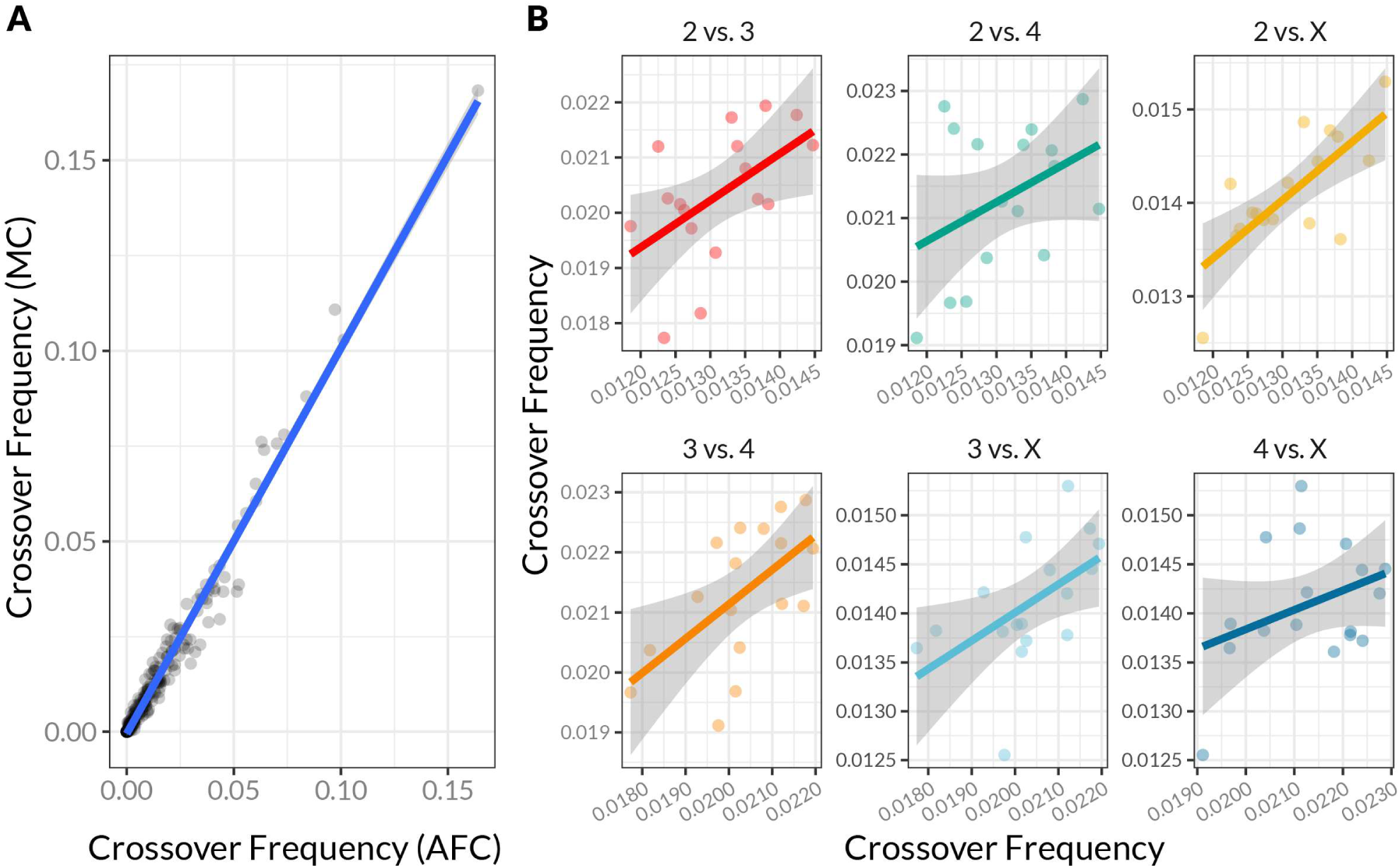
Recombination rate varies primarily at the genome-wide scale. (A) The correlation between recombination rate measured in genomic windows (∼300kb in size) in the MC and AFC populations. Each dot depicts a single genomic window (all chromosomes combined). (B) The correlation between chromosome-wide mean recombination rate between all pairs of chromosomes. Each point represents the recombination rate on two chromosomes for a single inbred line. Points and lines are colored to indicate the particular pair of chromosomes being compared. Positive trends indicate that recombination rates are consistent across chromosomes within lines (i.e. they vary genome-wide, and not idiosyncratically across chromosomes).

### *Q*_*ST*_-*F*_*ST*_

As expected from previous studies, genetic divergence between Madera Canyon, AZ and American Fork Canyon, UT was very low: genome-wide Weir and Cockerham’s *F*_*ST*_ was approximately 0.0039 (Figure 4A, mean *F*_*ST*_ of 6 591 high quality SNPs, MAF > 0.1, LD > 0.2). Examining variation in recombination, we estimated a within-population (between line) variance component of 0.066 and a between population variance component of 0.018, yielding na observed *Q*_*ST*_ of 0.212 (Figure 4A, dashed arrow). Our parametric bootstrap simulations of 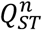 suggest that this value of *Q*_*ST*_ is highly unlikely to be observed under neutrality (0 of 10,000 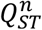 replicates were greater than the observed value of *Q*_*ST*_, thus p < 1×10^−6^). Similarly, the parametric bootstrap estimates of *Q*_*ST*_-*F*_*ST*_ under neutrality do not overlap the parametric bootstrap observed values of *Q*_*ST*_-*F*_*ST*_, even when taking into account sampling variance (Figure 3B). Together, these results indicate that while the observed phenotypic difference in recombination rate between MC and AFC is modest, it greatly exceeds its expected value under neutrality. This result is consistent with the hypothesis that natural selection underlies the observed difference in recombination rates between populations.

**Figure 4.**
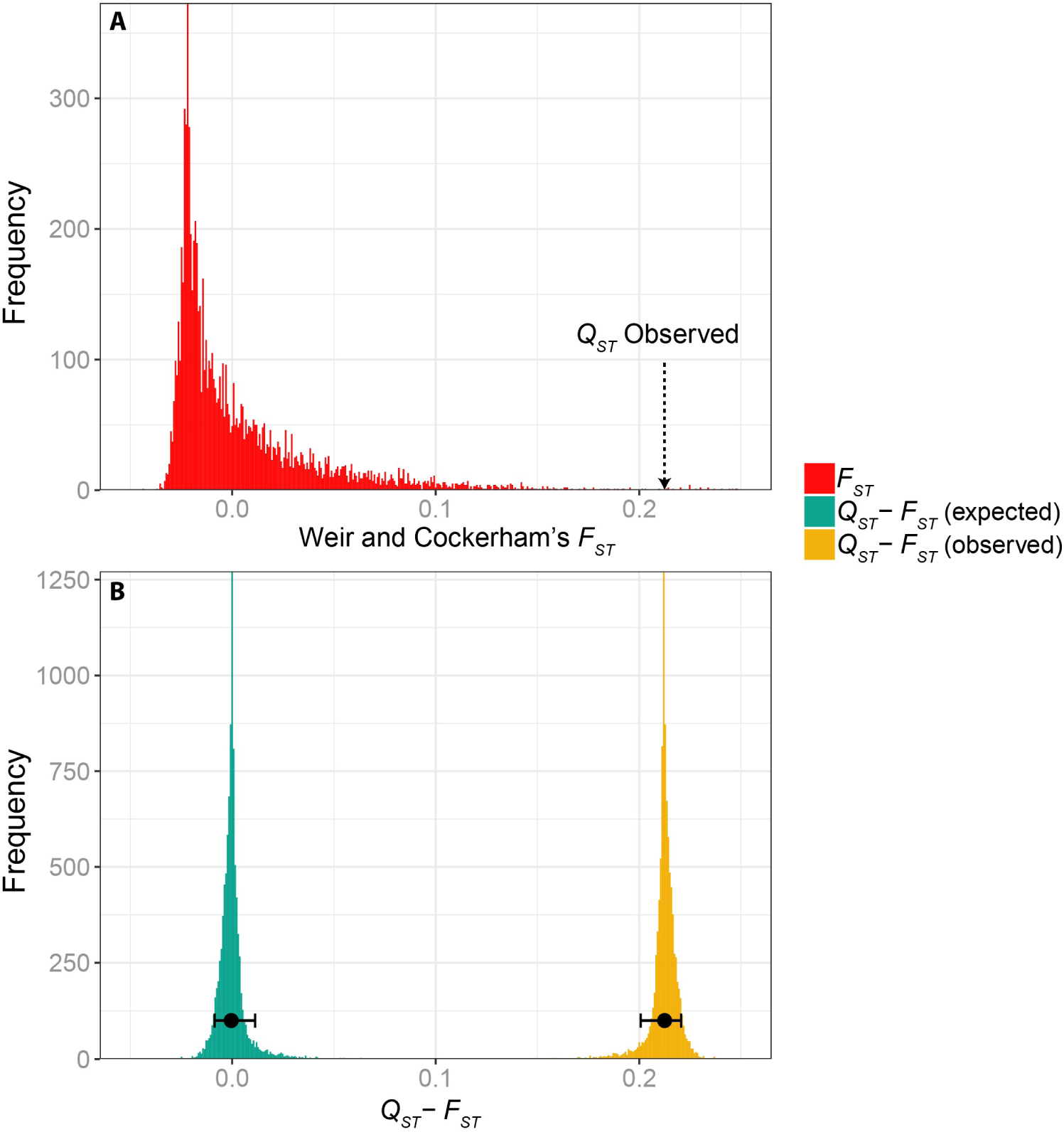
The value of Q_ST_-F_ST_ for recombination rate is greater than expected under neutrality. (A) Weir and Cockerham’s F_ST_ from 6591 RADseq-derived SNPs (mean F_ST_ = 0.0039). The observed value of Q_ST_ for recombination rate (0.212) is indicated with an arrow. (B) Comparisons of the sampling distribution of Q_ST_-F_ST_ expected under neutrality (green histogram) and the observed value (yellow histogram). Both distributions were simulated via a parametric bootstrap (see text). Black points with error bars indicate the mean and 95% confidence interval of the sampling distributions.

### Candidate genes

Of the 40 candidate genes examined, 33 had at least one non-synonymous polymorphism. Of these 33 genes, there were a total of 357 codons (out of a total of 29 964) with at least one non-synonymous polymorphism. After controlling for multiple comparisons three of these sites in two genes (*asp* and *mei-41*) were significantly associated with crossover rate (FDR adjusted p-value < 0.05, Figure 5A). Homozygous, non-reference, nonsynonymous changes at these three sites were variously associated with a 5%-7% differences in recombination between lines (Figure 5B). There was, however, strong LD (r^2^ > 0.8) between these alleles (e.g. lines with the lowest averaged crossover rates shared genotypic states for all three genes), and thus disentangling their independent effects on recombination rate was not possible.

**Figure 5.**
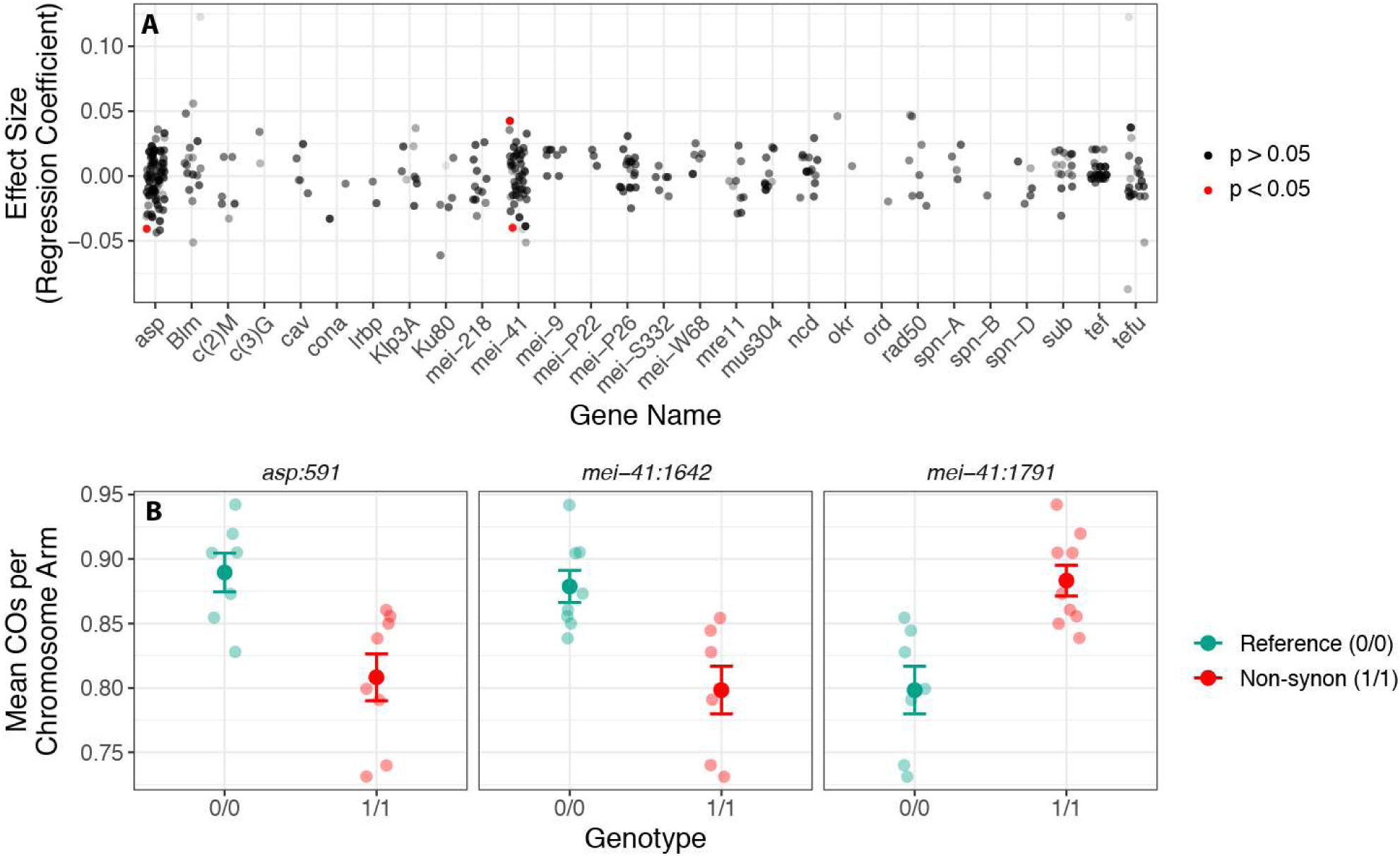
Non-synonymous substitutions in two meiosis-related genes are associated with variation in recombination rate. (A) Regression coefficients from linear models (y-axis) comparing genotype and crossover rate for sites (points) bearing non-synonymous, non-reference polymorphisms in a collection of meiosis-related candidate genes (x-axis). Red points indicate associations that were significant after adjustment via FDR correction (adjusted p < 0.05). (B) Mean recombination rates (crossovers per chromosome arm) for sites with significant associations (red points in A). Each panel depicts the mean and 95% confidence interval for crossover rates for each genotypic class (either homozygous reference or homozygous non-synonymous derived).

### Structural variation

Both short and long-read sequencing revealed extensive structural variation between inbred lines of *D*. *pseudoobscura*. As expected, the three strategies we used to detect structural variation (GATK INDELs, PacBio SV and LUMPY/Smoove) varied in the number and relative proportions of the various classes of structural variant they identified (Figure S3). That said, all three methods suggested that the most common form of structural variation are small to mid-sized INDELs, with larger deletions, insertions, and duplications being much rarer (Figure S2A). Consistent with the observation that AFC and MC are highly similar in their chromosomal arrangements, our structural variant analysis found no evidence of large-scale chromosomal inversions differentiating any of the lines.

Structural variation between lines did not co-vary with recombination rate (Figure 6). First, there was no relationship between recombination rate and the estimated percent sequence homology between the tester and inbred lines (Figure 6B, likelihood ratio test comparison of GLMMs, likelihood ratio test comparison of GLMMs *χ*^2^ = 2.9533, df = 3, p = 0.3989). Second, there was no relationship between recombination rate and the count of differences in structural variation between each inbred line and the tester line (Figure 6A, likelihood ratio test comparison of GLMMs *χ*^2^ = 1.1637, df = 3, p = 0.7617). This result was consistent across all methods used to identify structural variation (likelihood ratio tests, comparison of GLMMs with and without method by count/homology interaction effects, all p > 0.3). As such, at the 300kb scale, there is no evidence that the local differences in recombination rate among inbred lines are a result of differences in homology or local genome structure.

**Figure 6.**
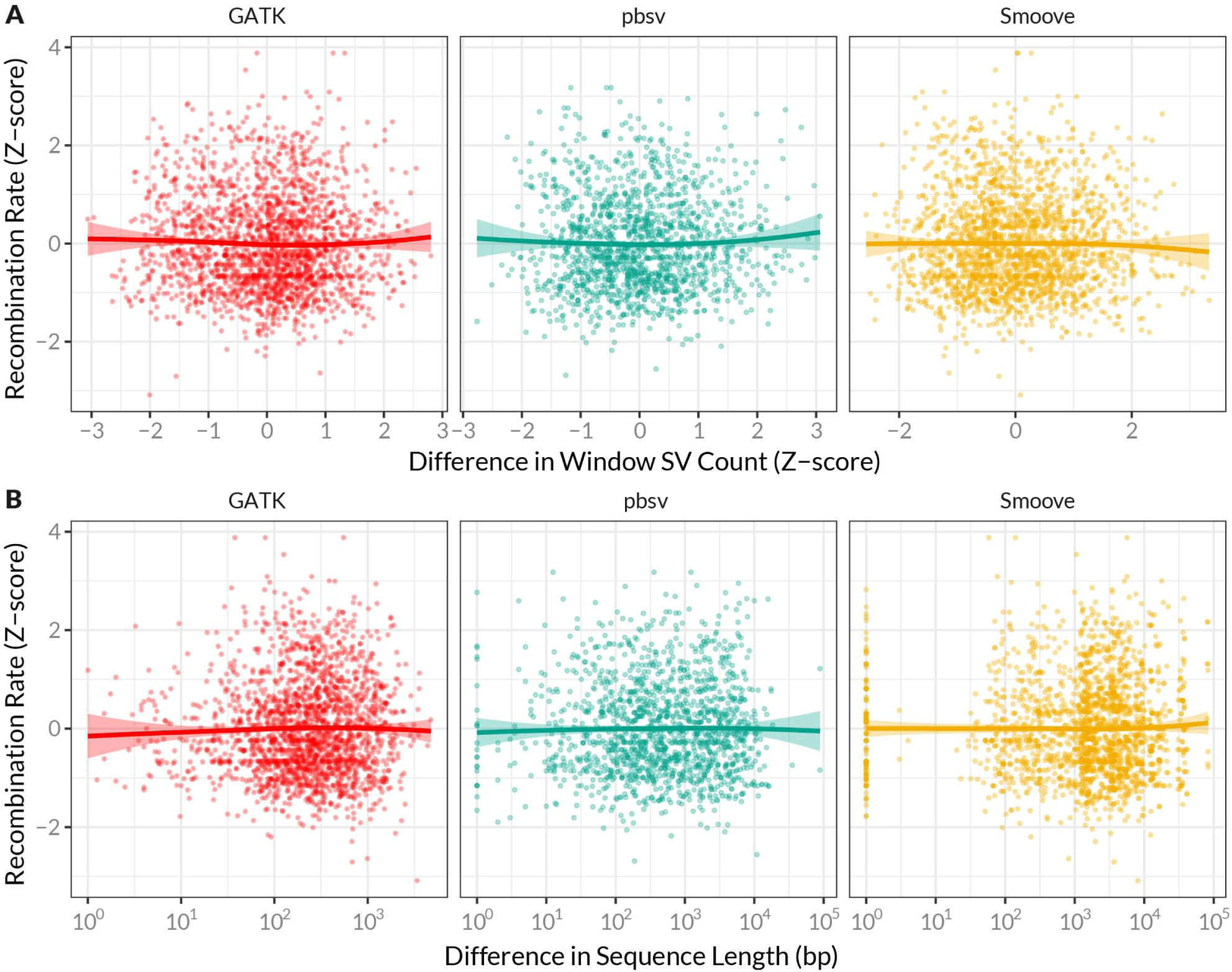
Structural variation is not correlated with recombination rate at the 300kb scale. (A) The relationship between normalized recombination rate and the normalized count of structural differences between each inbred line and the tester line. Each point represents a single recombination interval (all approximately 300kb in length) from one inbred line. Lines on each plot represent smoothed conditional means and are accompanied by 95% confidence intervals. Each column depicts the relationship using each of the three methods used to assay structural variation. (B) The relationship between normalized recombination rate and the difference in total sequence length between each inbred line and the tester line.

## Discussion

Recombination rate is a key modulator of many evolutionary processes, yet we have a poor understanding of how recombination rate itself evolves. Here, we studied how recombination rate varies using strains from two natural populations of *D*. *pseudoobscura* from Madera Canyon, AZ and American Fork Canyon, UT. We directly measured recombination rate in a total of 18 inbred lines from these populations and found substantial variation for recombination rate both within and between populations. Interestingly, the population from Madera Canyon, AZ exhibited an ∼8% lower recombination rate on average than the population from American Fork Canyon, UT. Within and between-population variation in recombination rate manifested largely as differences in genome-wide recombination rate, rather than changes in the local recombination landscape. This finding is supported by a general pattern of covariation in recombination rate among chromosomes within lines. While differences in recombination rates between populations were modest in absolute terms, a Q_ST_-F_ST_ analysis revealed that this difference vastly exceeds phenotypic divergence expected under neutral drift. This result is consistent with the hypothesis that local adaptation has driven differences in recombination rate between these populations.

We explored two possible mechanisms underlying recombination rate differences between lines. First, we found evidence that some differences in recombination rate between lines may involve non-synonymous coding changes in meiosis-related genes. Secondly, we found that local variation in recombination rate between lines does not correlate with local structural variation. These findings suggest that the differences in recombination we observed were driven by alleles resulting in genome-wide changes in recombination rate rather than local remodeling of the recombination landscape. Below, we discuss the relevance of our findings for the study of the evolution of recombination rate and relationships to previous work.

### Recombination rate variation in natural populations

Previous work has shown that recombination can vary between individuals (e.g. between crop varieties), or between species (Bauer *et al*., 2013; Stapley *et al*., 2017). However, to our knowledge, our study is the first to examine variation in recombination rate within and between natural populations of a single species. Our work demonstrates that there is ample genetic variation for recombination rate segregating in natural populations. As such, natural selection can likely readily act on recombination rate, even within a species.

Secondly, we found that variation in recombination rate was largely the result of variation in genome-wide crossover rates rather than changes in the recombination landscape. This suggests that changes in *genome-wide* recombination rate may be the predominant mechanism by which recombination rate evolves. This is particularly interesting given that many models of recombination evolution focus on the evolution of “modifier” alleles that alter genome wide rates of recombination (Barton & Otto, 1997; Otto & Lenormand, 2000). While this has been a useful simplification, our results suggest that true modifier alleles may exist to some degree in natural populations. Interestingly, recent work has shown that allelic variation in the coding sequence of the synaptonemal complex is correlated with variation in genome-wide recombination, providing a tantalizing molecular mechanism for modifier alleles (Wang *et al*., 2019).

To our knowledge, there is no *a priori* reason to expect recombination rate to vary predominantly at the genome-wide level. Indeed, there are reasons to expect the opposite. For one, the mutational target size of all possible variations in the recombination landscape (e.g., insertion or deletion of recombination-associated motifs) is likely much, much larger than all possible mutations of global regulators of recombination (e.g., genes in the meiosis pathway, (Ratnappan *et al*., 2012; Adrian *et al*., 2016; Howie *et al*., 2019). Given this, why did we observe largely global variation? One key consideration when interpreting this pattern is the effect of scale. While our inbred line sequencing data has effectively single base-pair resolution, the genetic markers we selected for our mapping populations were evenly distributed at approximately 300kb intervals. As such, our study was not designed to resolve differences in the fine-scale landscape of recombination. Such resolution would require sequencing substantially more markers and individuals per line in a direct trade-off with number of lines/populations (precluding the use of e.g. Q_ST_-F_ST_ type approaches). Apart from the effect of scale, another possibility is that there are indeed greater constraints on local rates of recombination than global rates, even at the 300kb scale. An example of this general phenomenon comes from systems with PRDM9-mediated recombination, in which PRDM9 binding motifs are strongly biased towards the upstream region of genes (Stevison *et al*., 2016; Baker *et al*., 2017; Paigen & Petkov, 2018). This bias ostensibly occurs to mitigate the mutagenic effects of recombination (Baker *et al*., 2017). It is possible that a similar process is operating at some scale in *Drosophila*, constraining the evolution of the recombination landscape locally, but not globally. Further work on local regulation of recombination may shed light on these constraints.

### Local adaptation of recombination rate

Our Q_ST_-F_ST_ analysis suggests that differences in recombination rate between *Drosophila pseudoobscura* populations from AZ and UT may have been driven by natural selection. To our knowledge, this is the first application of the Q_ST_-F_ST_ method to the study of recombination, and among the first evidence for the role of selection acting on genome-wide recombination rate in natural populations (Brand *et al*., 2018). However, while our results suggest a role for natural selection, the agent of selection underlying this change remains unknown. There are a wide variety of possible explanations for this difference (Dapper & Payseur, 2017). For example, differences in recombination between the populations may be directly favored, or other phenotypic differences may be divergently selected between the populations that incidentally affect recombination rate (via linkage or pleiotropy). One intriguing possibility is local differences in climate: recombination rate in *Drosophila* is known to be plastic with respect to ambient temperature (Jackson *et al*., 2015). Madera Canyon, Arizona has a mean annual temperature of approximately eleven degrees Celsius higher than American Fork Canyon, Utah (10.5C vs 21.6C, (Menne *et al*., 2018). This increase in temperature likely causes a facultative increase in recombination rate in the Madera Canyon population. We speculate that the difference in recombination rate we observed under constant conditions may be a compensatory response to an environmentally-induced increase in recombination rates, in order to return genome-wide recombination rate to some optimum value (i.e., a response to maladaptive plasticity, King & Hadfield, 2019). That said, further work will be needed to connect variation in recombination rates to specific agents of selection. One obvious extension of our approach would be a greater number of populations, perhaps existing over a climatic gradient (or paired populations in differing environments). We hope that our demonstration of the efficacy of the Q_ST_-F_ST_ method inspires the undertaking of such eco-evolutionary studies of recombination rate.

### Structural variation as a modulator of recombination rate

We found no association between among-line variation in recombination rate and among-line variation in the abundance or size of structural variants. An important consideration here is that that this analysis was not intended to test whether *average* recombination rate (across all lines) is associated with structural variation – this association is extremely well documented and is unquestionably present in our data (Ross *et al*., 2015; Kent *et al*., 2017; Mugal *et al*., 2017). Instead, our goal was to test if *among-line variation* in recombination rate in each genomic interval was explained by among-line structural differences, using normalized metrics of both recombination rate and structural variation within each genomic interval (as Z-scores, i.e. statistical controlling for average recombination rate).

Why was there no detectable association between structural variation and local rates of recombination? The scale of our recombination estimates is also important to consider here: much of the previous work describing the effects of heterozygous structural variation on crossing-over was performed at very fine scale, e.g. <1kb in Arabidopsis (Opperman *et al*., 2004). It may be that changes in recombination resulting from structural variation are restricted to finer genomic scales (i.e. <300kb) and that other types of regulators (e.g. variation in meiosis genes or the chromatin landscape) modulate recombination at larger scale (Brand *et al*., 2018). A notable exception to this is large scale chromosomal inversions (notably absent in our lines), which are well known to affect recombination at scales much larger than 300kb – upwards of 10Mb in many cases (Crown *et al*., 2018; Said *et al*., 2018). However, inversions likely have outsized recombination suppressing effects compared to other forms of non-homology because of the loop structures they form during chromosome pairing (Ortiz-Barrientos *et al*., 2006; Crown *et al*., 2018). Further work will be required to disentangle the relative contribution of structural and global/trans modifiers of recombination rate at different genomic scales.

### Amplicon sequencing as a tool for genetic maps

Our ability to economically sequence hundreds of markers in thousands of individuals was made possible by the GT-seq amplicon sequencing approach (Campbell *et al*., 2015). This technique is highly scalable, and in our case, we likely could have sequenced many more markers (and/or individuals) while maintaining a very high depth per amplicon. This method is an alternative to the increasingly popular bulk-sequencing approaches, in which sample DNA is pooled prior to sequencing (Chakraborty *et al*., 2018). GT-seq avoids some of the complexity of these approaches. For one, because it is a PCR-based method, GT-seq does not require performing extraction, quantification and manual normalization of sample DNA. This is a non-trivial consideration when individual sample sizes are in the thousands. Further, unlike bulk-sequencing, amplicon sequencing provides individual-level genotypes. As such, the occurrence of double/triple/etc. crossovers can be directly resolved, and problematic individuals identified and removed during analyses. To our knowledge, these are both not currently possible with bulk sequencing (unless barcodes are employed, limiting the total number of individuals in the pool). The main drawbacks of amplicon sequencing are a decrease in resolution (number of markers), and the need to pre-identify mapping informative markers. That said, we believe GT-seq and amplicon sequencing more generally will be a useful tool for future studies of variation in recombination rate and can be readily paired with other approaches depending on the goals of the study.

### Conclusion

Recombination rate plays and important modulatory role in many evolutionary processes, but little is known about how recombination rate itself evolves. Here, we studied natural variation in recombination rate within and between two populations of *Drosophila pseudoobscura*. We found extensive genetic variation for recombination rate within and between populations, with the majority of variation detected manifesting as differences in overall genome-wide recombination rate. This suggests that the differences in recombination we detected between lines may be the result of genetic variation in trans-acting global regulators of recombination, an idea supported by a significant association between non-synonymous variation in meiosis-associated genes and recombination rate. We also found no evidence that among-line differences in local recombination rate at the 300kb scale were correlated with structural variation within the lines. Finally, we discovered that the magnitude of phenotypic difference in recombination rate between the two populations was far greater than expected under a model of neutral trait evolution, suggesting that the differences may have been driven by natural selection. Our study provides key insights in the quantitative genetics of recombination rate and lays the groundwork for future research focused on studying the recombination rate in natural populations.

## Acknowledgements

Funding for this project was provided by National Science Foundation grants DEB-1545627, 1754022, and 1754439 to MAFN. KMS was additionally supported by a Natural Sciences and Engineering Research Council of Canada Postdoctoral Fellowship. The tester line (MV2-25) was generously provided by Dr. Steven Schaeffer. We thank Dr. Noah Whiteman and Dr. John Chaston for assistance during fieldwork. Dr. Katharine Korunes, Dr. Jenn Coughlan and members of the Noor Lab provided valuable feedback on the manuscript. Dr. Andrew MacDonald provided guidance on statistical analyses. Dr. Armin Töpfer provided advice on the use of the pbsv software.

