## Supplemental Figures for "Natural selection shapes variation in genome-wide recombination rate in *Drosophila pseudoobscura*"

**SUPPLEMENTARY FIGURES**

The evolutionary genetics of recombination rate in natural populations of *Drosophila pseudoobscura*

*Samuk et al. 2019*

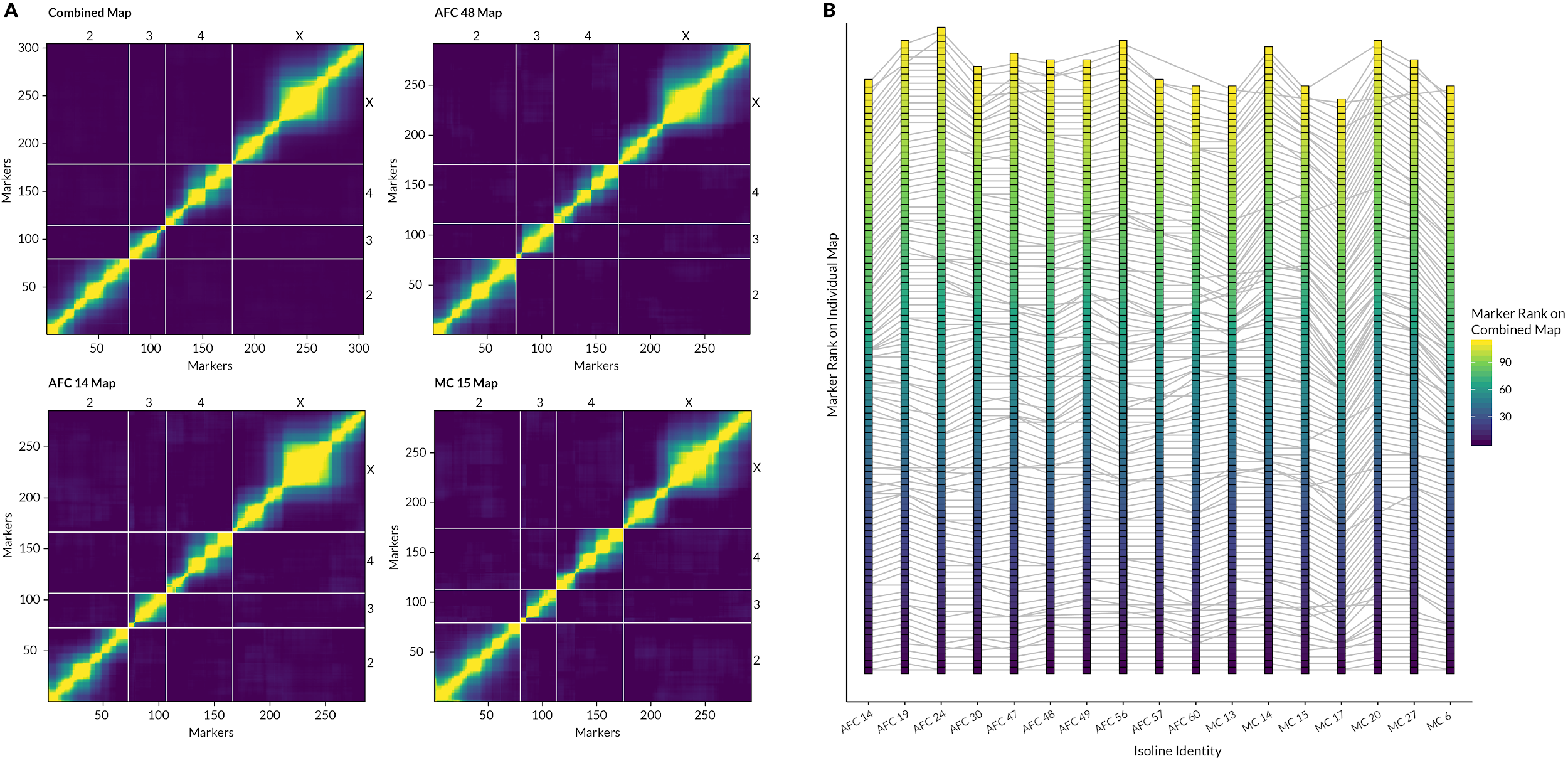

**Figure S1 |** Comparison of inferred marker orders in combined vs. individual datasets. (A) Recombination fraction heat maps for four representative maps. The combined map was built by pooling all samples before inferring marker order. Lighter colors correspond to larger recombination fractions between pairs of markers. (B) Inferred markers orders on Chromosome X for all isolines. Each square point is a single marker on chromosome X, and markers are ordered based on their rank (y-axis) in each isoline-level individual map (x-axis).horizontal lines between points show the position of the same marker in each individual map. The color scale indicates the rank inferred from the combined map: major changes in marker order between the combined and individual maps would manifest as jumbled colors in the gradient, of which there are none.

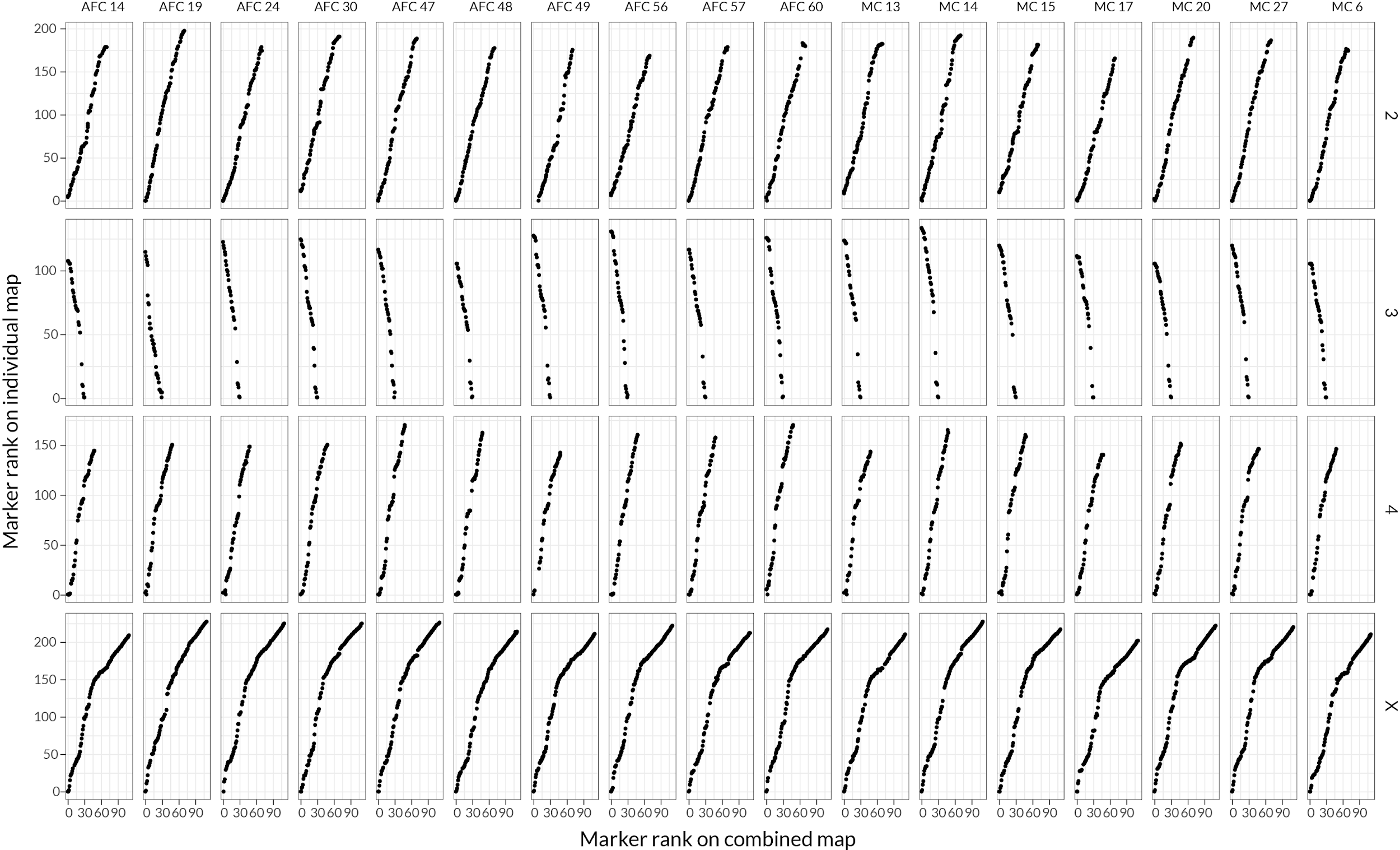

**Figure S2 |** Genome-wide comparison of inferred marker orders in combined vs. individual datasets. Each plot displays the inferred rank order marker for markers in the individual map (y-axis) and the combined map (x-axis) for a single chromosome in a single isoline. Preservation of marker order is reflected by a progressive increase in marker rank on both axes, with no changes in directionality (e.g. due to inversions or misordered markers).

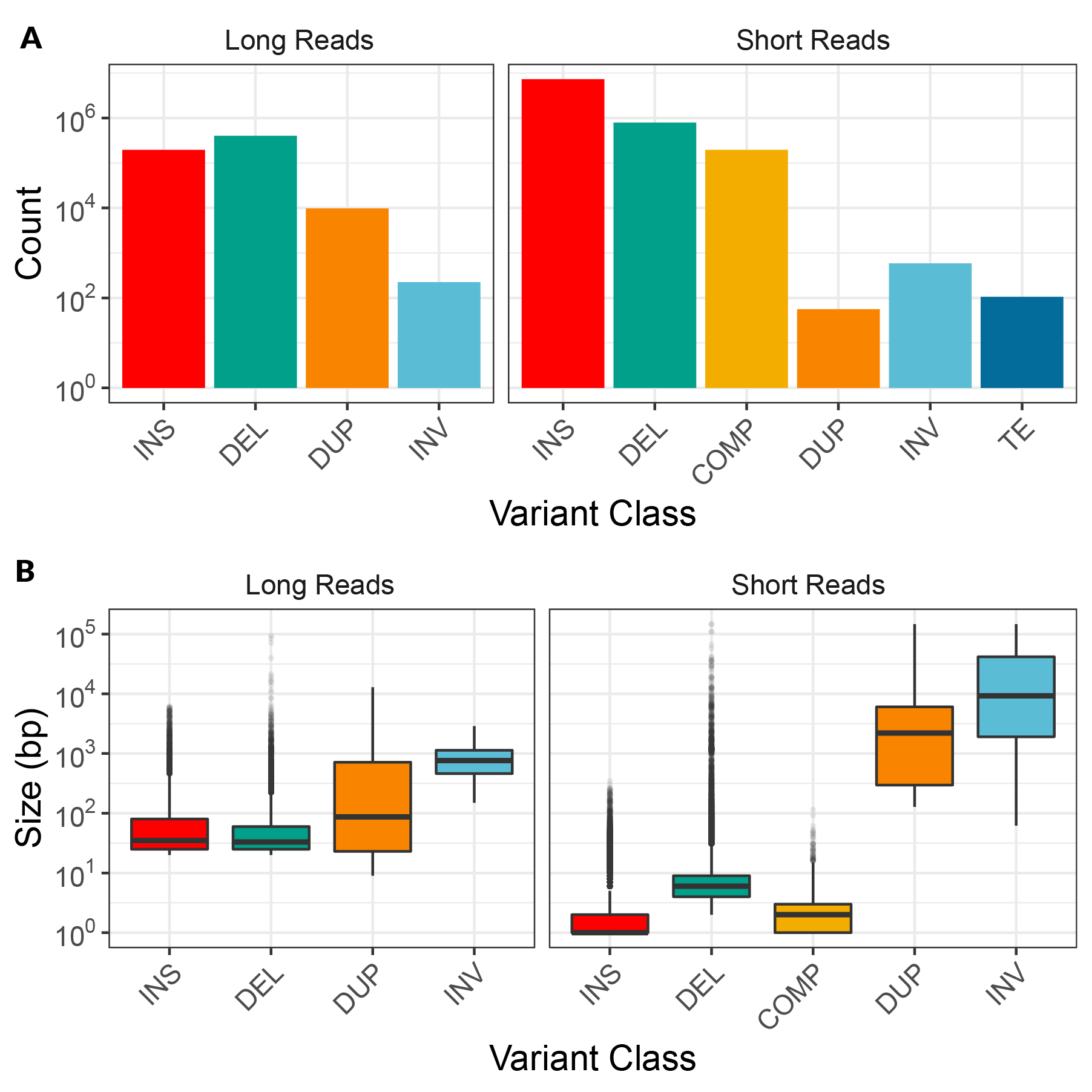

**Figure S3|** Summary of the count (A) and size in base pairs (B) of structural variants identified via short and long read sequencing. Structural variant classes are: insertions (INS), deletions (DEL), complex indels (COMP), duplications (DUP), inversions (INV), transposable elements (TE).
